# Early life adversity drives sex-dependent changes in 5-mC DNA methylation of parvalbumin cells in the prefrontal cortex in rats

**DOI:** 10.1101/2024.01.31.578313

**Authors:** Emma S. Noel, Alissa Chen, Yanevith A. Peña, Jennifer A. Honeycutt

## Abstract

Early life adversity (ELA) can result in increased risk for developing affective disorders, such as anxiety or depression, later in life, with women showing increased risk. Interactions between an individual’s genes and their environment play key roles in producing, as well as mitigating, later life neuropathology. Our current understanding of the underlying epigenomic drivers of ELA associated anxiety and depression are limited, and this stems in part from the complexity of underlying biochemical processes associated with how early experiences shapes later life behavior. Epigenetic alterations, or experience-driven modifications to DNA, can be leveraged to understand the interplay between genes and the environment. The present study characterized DNA methylation patterning, assessed via evaluation of 5-methylcytosine (5-mC), following ELA in a Sprague Dawley rat model of ELA induced by early caregiver deprivation. This study utilized maternal separation to investigate sex- and age-specific outcomes of ELA on epigenetic patterning in parvalbumin (PV)-containing interneurons in the prefrontal cortex (PFC), a subpopulation of inhibitory neurons which are associated with ELA and affective dysfunction. While global analysis of 5-mC methylation and CpG site specific pyrosequencing of the PV promoter, Pvalb, showed no obvious effects of ELA, when analyses were restricted to assessing 5-mC intensity in colocalized PV cells, there were significant sex and age dependent effects. We found that ELA leads sex-specific changes in PV cell counts, and that cell counts can be predicted by 5-mC intensity, with males and females showing distinct patterns of methylation and PV outcomes. ELA also produced sex-specific effects in corticosterone reactivity, with juvenile females showing a blunted stress hormone response compared to controls. Overall, ELA led to a sex-specific developmental shift in PV profile, which is comparable to profiles that are seen at a later developmental timepoint, and this shift may be mediated in part by epigenomic alterations driven by altered DNA methylation.

## INTRODUCTION

Early life adversity (ELA) can increase susceptibility to developing anxiety disorders, many of which manifest in a protracted manner (McEwen, 2017), and has profound impacts on overall physiological and mental health (Luby et al., 2017). While it has long been documented and accepted that ELA is a significant risk factor for the development of affective disorders (i.e., anxiety, depression) (Bandelow et al., 2004; Kaplow and Windom, 2007; Copeland et al., 2018, Targum and Nemeroff, 2019), the neurobiological and epigenomic changes driving risk remain to be fully understood. Brain development is dictated by interactions between genetic predisposition and environmental factors, which drive individual differences in neuropathology and behavioral responses acutely and later in life (Kolb and Gibb, 2011). Notably, women are at higher risk for developing ELA-associated disorders like anxiety, depression, and posttraumatic stress disorder (PTSD) (Kessler, 1994; Kessler et al., 1996; Veijola et al., 1998; Tolin and Foa, 2008; Albert, 2015), though they remain underrepresented in the literature (McCarthy et al., 2012; Shansky, 2019). Adverse experiences during early life, when the brain is most plastic and sensitive to environmental and experiential influences, can significantly alter neuronal development (Galuske et al., 2019), leading to epigenetically-driven changes in affective processing associated with emotional disorders (Luoni et al., 2015). Indeed, emerging research has begun to identify epigenetic changes in genes linked to ELA associated risk and mental illness (e.g., Palma-Gudiel et al., 2015; Non et al., 2016; Fachim et al., 2021), findings which will no doubt pave the way for targeted intervention and/or treatment. However, we are only truly beginning to understand the complexity with which early environment confers changes to the epigenome, and how those changes influence neural pathology and concomitant behavior.

While short-term exposure to adverse experiences may produce resilience (McGloin and Widon, 2001; Collishaw et al., 2007), the manifestation of anxiety-related disorders can arise from chronic adversities, particularly those experienced in childhood (Green et al., 2010; Nanni et al., 2012; McEwan, 2017). Moreover, hyper-activation within the stress response system can impact stress-related hormones (i.e., cortisol in humans or corticosterone in rodents) (Kalinichev et al., 2022; Alexander et al., 2018), and has been shown to reduce gene expression associated with neuroplasticity (Begni et al., 2020). The prefrontal cortex (PFC) is particularly vulnerable to the impact of ELA due to its protracted development, providing an ample developmental window within which adverse experiences can impart epigenetic modifications (Kundakovic and Champagne, 2014). Indeed, childhood ELA alters neurodevelopment of the PFC and its connections with other brain regions, with mounting evidence suggesting that ELA drives precocial maturation of corticolimbic circuitry in both humans (e.g., Gee et al., 2013) and in rodent models (e.g., Honeycutt et al., 2020), and likely contributes to maladaptive behaviors associated with affective dysfunction. PFC function regulates responses to possible threat in the environment, and disruption can lead to anxiety-like behaviors in preclinical models of ELA that differ based on sex, though the directionality of these sex differences is unclear as so few studies leveraging ELA models have systematically examined sex as a biological variable (Ellis and Honeycutt, 2021).

The PFC contains two subregions – the prelimbic (PL) and infralimbic (IL) – that are linked to affective processing, threat response, and inhibitory control (Capuzzo and Floresco, 2020; Kredlow et al., 2022). The PL PFC is involved in processing higher-order cues and adapting behavioral response to changing environmental stimuli (Sharpe and Killcross, 2015), while the infralimbic (IL) region is implicated in stress response (Pace et al. 2020). Together, these brain regions work to respond to aversive stimuli (Giustino and Maren, 2015), with the IL showing changes in local excitatory: inhibitory (E:I) balance following chronic stress (Pace et al., 2020). Additionally, the PFC and basolateral amygdala (BLA), a region known for its involvement in fear (Sharp, 2017), emotionality (Terburg et al., 2012), and anxiety (Prager et al., 2016), together coordinate responses to stressors (Stevenson et al., 2008) and both regions are implicated in the emergence and persistence of anxiety-like responses following ELA (Giustino and Maren, 2015; Prager et al., 2016). Within the PFC, tightly controlled E:I balance is necessary for essential aspects of decision making, working memory and overall executive functioning (Ferguson and Gao, 2018). Disruption of the E:I balance has been documented in patients with anxiety disorders, with increased activity of the PFC reported (Martin et al., 2009).

A primary regulator of E:I balance in the PFC are a subset of inhibitory GABAergic neurons that express the calcium-binding protein parvalbumin (PV; Ferguson and Gao, 2018), and is encoded by the Pvalb gene. PV-expressing neurons are fast-spiking and provide perisomatic inhibition that positions them to efficiently regulate local tone and orchestrate neuronal firing and synchrony (Whittington et al., 2011). As such, they are implicated in many dysregulated pathways related to neuropsychiatric disorders (Mannekote Thippaiah et al., 2022). PV is also a critical factor in regulating calcium-dependent processes as a slow calcium buffer that helps synchronize short-term synaptic plasticity, making PV essential in neuroplasticity and behavior (Caillard et al., 2000; Permyakov and Uversky, 2022). Moreover, reductions in PV are linked to alterations in GABA release, including asynchrony in neuronal firing (Permyakov and Uversky, 2022; Collin et al., 2005). Notably, research has shown that ELA confers changes in the number of PV neurons in the PFC (Hu et al., 2010a; Lussier and Stevens, 2016; Perlman et al., 2021), which may also be sex-specific (Ellis and Honeycutt, 2021). This has led researchers to hypothesize that decreased PV protein expression may contribute to alterations in PV cell count, expression, and/or function, thereby leading to deleterious outcomes. In PV+ knockout mice, reduced social behavior was observed in response to lowered PV expression (Wöhr et al., 2015). This study suggests down-regulation in the transcription or translation of the protein itself is a key contributing factor to altered behavior, not necessarily of the loss of PV-containing interneurons (though both may occur at the same time). This distinction is important, as it suggests that PV serves not only as a molecular marker of a subset of interneurons, but also as a potential regulator of behavior. PV expression also changes following ELA (Honeycutt and Ellis, 2021) and in neuropsychiatric disorders including schizophrenia, anxiety, and depression, making PV a prospective player in the development of these disorders (Ferguson and Gao, 2018).

ELA is generally thought to produce a decrease in PV expression in the PFC (Brenhouse and Andersen, 2011; Ellis and Honeycutt, 2021; Page et al., 2019; Perlman et al., 2021; Zaletel et al., 2016). ELA also appears to induce sex-specific differences in PV expression, as female rats show a decrease in PV protein levels specifically in the PFC during juvenility (Holland et al., 2014). Additionally, ELA-exposed females show earlier maturation of BLA-PFC circuitry, and earlier onset of anxiety-like behavior, both of which are sustained at the functional level across development (Honeycutt et al., 2020). This juvenile female specific maturation pattern is potentially associated with changes in PV levels and/or function. In conjunction with clinical studies, women show earlier emergence of anxiety following ELA than men, suggestive of earlier underlying neurobiological changes (Goodwill et al., 2019). Altogether, PV has emerged as a promising target of investigation in the sex-specific manifestation of mood disorders and may be driven by epigenomic changes conferred from early life experiences.

ELA leads to epigenetic changes that can alter gene expression, leading to stable changes to the epigenome (Anacker et al., 2014; Dammann et al., 2011; Dong et al., 2015; Jawahar et al., 2015; Szyf, 2013a). DNA methylation is an epigenetic modification of interest and is the covalent linkage of a methyl group onto a cytosine base pair to create 5-methylcytosine (5-mC). DNA methylation can disrupt gene expression by interfering with transcription factor binding or by other downstream mechanisms (Szyf, 2013a). In both humans and animal models, alterations to DNA methylation patterns on specific CpG sites (e.g., Humphreys et al., 2019), and globally (e.g., McCoy et al., 2019), are associated with risk for anxiety and depression. Historically, changes in DNA methylation were thought to only be involved in cellular differentiation (Huang and Fan, 2010; Szyf, 2013a). More recently, DNA methylation has been identified as an epigenetic regulator of genes and thought to be a rapid, relatively stable epigenomic alteration that is highly regulated by neuronal activity (Guo et al., 2011; Kim and Costello, 2017). Methylation in the promoter region of a gene can interfere with transcription factor binding, and therefore alter gene regulation and subsequent protein production (Héberlé and Bardet, 2019). Thus, DNA methylation is implicated in the silencing of genes following impactful early life experiences.

A number of DNA methylation studies pertaining to ELA have attempted to find a common linkage between ELA, DNA methylation sites and directionality of methylation (i.e., hypo- or hyper-), with varying results across species, brain region, and other treatment and/or experiential conditions (e.g., Czamara et al., 2021; Dammann et al., 2011; Daniels et al., 2009; Esposito et al., 2016; Fiacco et al., 2019; Fiori and Turecki, 2016; Fitzgerald et al., 2021). Some studies suggest that DNA methylation may serve as a biomarker for either resilience (Kundakovic et al., 2015) or vulnerability (Cattaneo and Riva, 2016) to the development of psychiatric disorders. In ELA exposed rats, reduced global DNA methylation in the amygdala was correlated with a decrease in anxiety-like and depressive-like behavior (McCoy et al., 2019), suggesting DNA methylation as a potential regulator of behavior. Indeed, evidence suggests that blockade of methylation early in life can prevent certain behavioral outcomes, such as anxiety (Weaver et al., 2004), further underscoring an intriguing relationship between these two factors. In humans, global increases in DNA methylation in the blood and PFC (Houtepen et al., 2016) have been associated with ELA (Kundakovic et al., 2015; Szyf, 2013a) which also corresponded with hyperactivity in HPA axis response (Oldehinkel and Bouma, 2011). However, gross changes in global DNA methylation cannot identify the loci of methylation action, therefore it is important to identify and characterize genes susceptible to ELA induced methylation.

Throughout development in rodents, PV cell count increases exponentially and then decreases as neural pruning occurs following juvenility (Ellis and Honeycutt, 2021). Decreases in PV expression (Gildawie et al., 2020) and cell count (Lussier and Stevens, 2016) can occur in response to ELA, and are implicated in the pathophysiology of psychiatric disorders (Ferguson and Gao, 2018). Decreases in PV expression are frequently documented in schizophrenia patients, and in rodent models of schizophrenia (Fachim et al., 2016; 2018; Lodge et al., 2009). Patients with schizophrenia show reduced PFC density of PV-containing interneurons (Kaar et al., 2019). Additionally, in humans with schizophrenia or major depressive disorder, and in mouse models of schizophrenia, hypermethylation of the Pvalb promoter is associated with decreased PV protein (Fachim et al., 2018; Thaweethee-Sukjai et al., 2019). Changes in PV protein output in response to methylation suggest that the Pvalb promoter region is sensitive to methylation, and that methylation impacts the transcriptional profile of PV. Therefore, DNA methylation related to PV cells is a promising target for investigation in the pathophysiology of ELA, and that changes to PV outcomes may be mediated by DNA methylation accumulation (e.g., 5-mC) at the promotor.

Overall, ELA contributes to changes in DNA methylation; however, the directionality of the relationship between DNA methylation and concomitant psychopathology remains unclear. Furthermore, methodological and end-point analyses make it difficult to identify specific methylation loci that drive the emergence of distinct pathological profiles. Thus, the present work addresses methylation profiles in global, cell-specific, and gene-specific assays. This will help paint a broader picture of how ELA alters the epigenome to impact gene expression, and if these changes endure throughout development in males and females. Sex-differences persist in anxiety disorders, and female rats show earlier maturation of PFC-BLA circuitry (Honeycutt et al., 2020) as well as disruptions in PV levels (Ellis and Honeycutt, 2021). Thus, this work focuses on the PFC, which is heavily implicated in anxiety response following ELA. Following ELA, PV cell counts, 5-mC signal intensity, global DNA methylation percentages, and colocalization of 5-mC with PV, along with anxiety-like behavior and physiological response via corticosterone reactivity were assessed. Overall, we found sex-specific across measures, suggesting a differential impact of ELA on PV and 5-mc outcomes. Following ELA, juvenile female rats displayed lowered PV+ cell count with higher intensity of 5-mC in PV+ cells, and lower corticosterone reactivity, whereas ELA males showed increased PV cell counts and reduced 5-mC intensity in PV cells, with no changes to corticosterone reactivity. Neural outcomes for both PV and 5-mC in juvenile ELA rats were comparable to those observed in young adults and suggest that ELA may differentially induce earlier maturation of epigenomic outcomes, which may alter PV function and increase risk for affective dysfunction.

## METHODS

### Subjects and Early Life Adversity via Maternal Separation

Timed-pregnant Sprague-Dawley rat dams (Charles River Labs, Wilmington, MA) arrived at gestational day (GD) 15 with parturition designated as postnatal day (P) 0. After birth (P0), whole litters were randomly assigned to either control (CON; standard rearing) or early life adversity (ELA; via maternal separation). From each litter, no more than two rats per condition were used to avoid possible litter effects. On P1, litters were culled to 10 (+/-2) with an equal ratio of males and females maintained wherever possible (Agnish, 1997). Following weaning (P21), all offspring were housed in same-sex pairs and left undisturbed until behavioral testing (either as P25 juveniles or P45 young adults), except for regular cage changes. All dams and offspring used for this study were kept in a temperature- and humidity-controlled vivarium with a 12-hour light cycle (lights on at 0700h) and had *ad libitum* access to food and water.

Litters assigned to ELA rearing conditions were subjected to daily maternal separation, which is a well-established model of caregiver deprivation (e.g., Coley et al., 2019; Ganguly et al., 2019; for review, see Nishi, 2020). From P2-20, ELA litters underwent maternal separation. From P2-10, pups were separated from dam and littermates and placed in individual cups with pine shavings from the home cage and thermoregulated in a circulated water bath (37°C) for 3.5 hours each day. From P11-20, when they can thermoregulate, pups were placed in individual cages with home cage pine shaving beddings for 4 hours per day. CON litters remained with dams from P2-20 but were individually handled on P11 and P15 for acclimation to experimenter interactions, in addition to periodic weighing experienced by all pups to track development. See Figure 1 for experimental timeline of rearing, behavior, and tissue collection endpoints. A total of 80 offspring were used for the present study (*n*=40 CON; *n*=40 ELA), which consisted of behavioral testing at 2 different ages (P25 or P45) followed by immunohistochemistry (IHC) for 5-mC and PV in one brain hemisphere, and DNA extractions occurring with the other.

**Figure 1.**
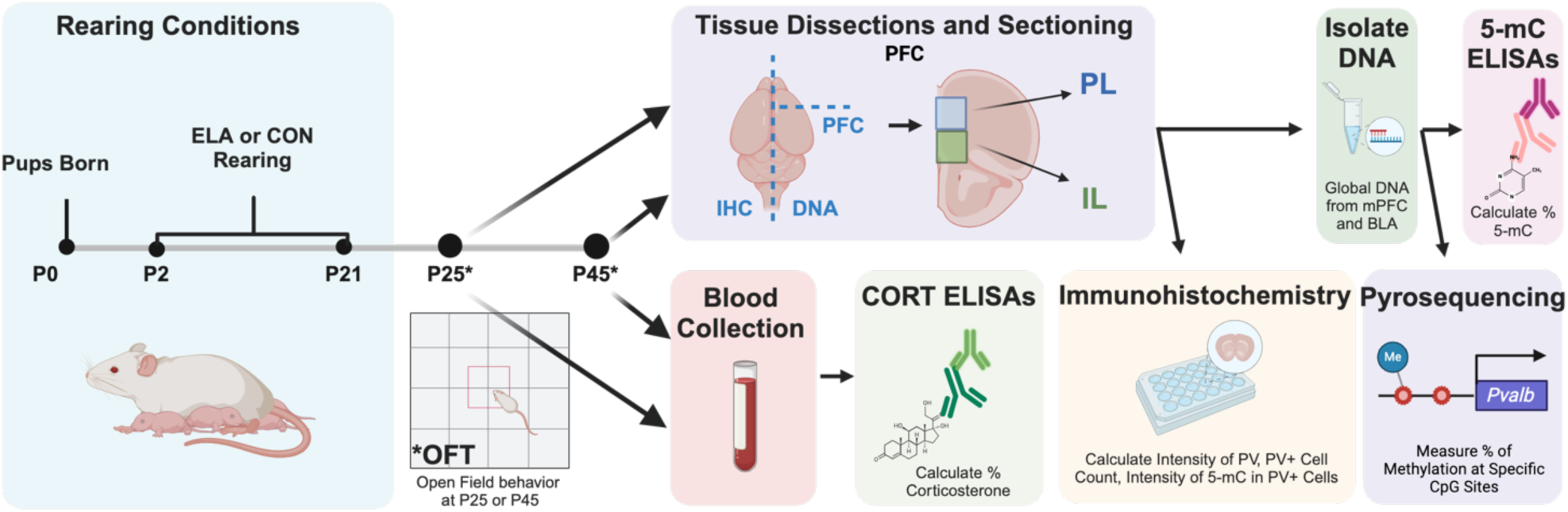
Experimental Timeline. Timed pregnant rat dams give birth (P0), and pups experience either CON or ELA rearing from P2-21, when they are weaned. At P25 or P45, rats are assessed for anxiety-like behavior in the OFT for 10 minutes. Following behavior, samples are collected following euthanasia. Corticosterone levels are assessed though collected trunk blood samples with an ELISA. Brains are extracted and bisected by hemisphere, with one hemisphere postfixed in 4% paraformaldehyde for latter IHC assessment of PV, 5-mC, and 5-mC intensity in colocalized PV cells. The other hemisphere is rapidly microdissected to isolate the PFC to extract purified DNA samples for 5-mC methylation ELISA and pyrosequencing assays. *Abbreviations*: P (postnatal day); CON (Control); ELA (Early Life Adversity); OFT (open field test); PFC (prefrontal cortex); PL (prelimbic PFC); IL (infralimbic PFC); IHC (immunohistochemistry); CORT (corticosterone); 5-mC (5-methylcytosine). Figure generated with BioRender.

### Behavioral Testing: Open Field Test (OFT)

All animals were assessed for anxiety-like behavior in the open field test (OFT). For two consecutive days prior to testing, rats were habituated for 30 minutes in the behavioral testing room in their home cages with cage mates. On testing day, rats were transported in a clean holding cage with pine shavings into a dimly lit testing room. Following behavioral testing, rats were returned to their home cage in the colony. The OFT was conducted in a 100cm x 100cm black plexiglass square arena with 40cm tall walls (Maze Engineers). Each rat began testing facing the same corner and were allowed given 10 minutes to explore the arena uninterrupted. After each test, the arena was cleaned with 50% ethanol solution to remove olfactory cues. A webcam mounted to the ceiling (Color GigE Camera, Ethovision) above the arena recorded the full session and videos archived. Recordings were analyzed for time (s) spent in center and distance traveled (cm) with Noldus Ethovision 16.0 XT software.

### Tissue and Blood Collection

Following completion of behavioral testing, rats were deeply anesthetized using CO2 and rapidly decapitated. Approximately 1.5mL of trunk blood was collected in individual tubes for later corticosterone assessment from each rat. Brains were immediately extracted, and one hemisphere was rapidly dissected on ice to isolate PFC samples (Figure 2A). Once isolated, PFC samples were placed into microcentrifuge tubes, flash frozen (in 100% ethanol surrounded by dry ice) and stored at -80°C for later DNA extractions. The second hemisphere was placed in 4% paraformaldehyde solution for 1 week of post fixation at 4°C for later immunohistochemistry (IHC) processing and microscopy analyses (see Figure 2B). Brain hemisphere allocation was pseudorandomized across all subjects/conditions, such that both hemispheres were included in methylation and IHC analyses.

**Figure 2.**
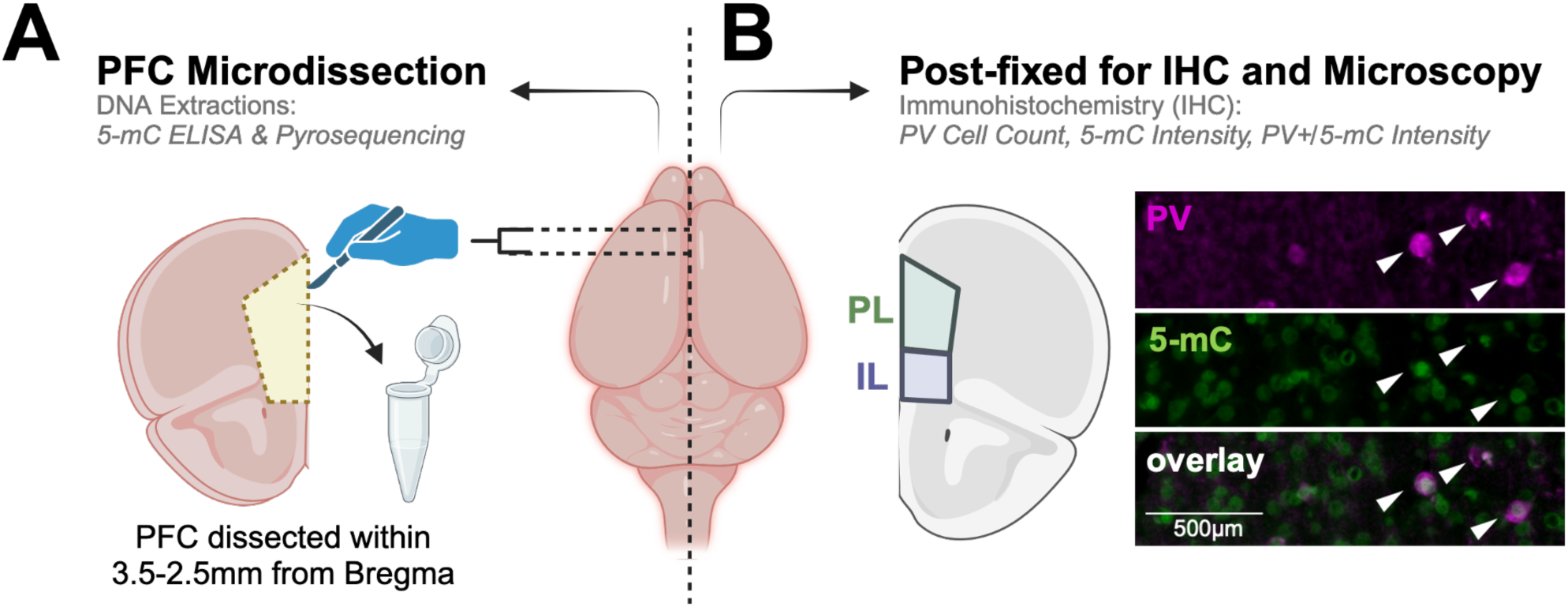
Neuroanatomical Isolation of PL and IL PFC Samples for DNA Extraction and Immunostaining Experiments. Upon extraction, brains were rapidly microdissected to isolate PFC tissue within +3.5 to +2.5mm relative to Bregma in one pseudorandomly selected hemisphere. Once isolated, whole PFC samples (containing both PL and IL regions) were snap frozen and stored at -80°C until tissue homogenization and DNA extraction procedures (**A**). Extracted DNA was used for in-house 5-mC DNA methylation ELISA and remainder was sent to an outside vendor for pyrosequencing. The remaining hemisphere was postfixed in 4% paraformaldehyde, sectioned, and immunostained for PV and 5-mC (**B**). Representative images of individual immunostained neuronal markers and merged image can be seen in (**B**). Microscopic imaging was conducted to visualize these markers in both the PL and IL PFC for later quantification. Abbreviations: PFC (prefrontal cortex); PL (prelimbic PFC); IL (infralimbic PFC); PV (parvalbumin); 5-mC (5-methylcytosine) Figure generated with BioRender.

### Corticosterone ELISA

Blood samples were left at room temperature for 1 hour to coagulate, and then spun down (1500rpm) in a microcentrifuge at 4°C for 10 minutes. Once separated, serum was collected and stored at -20°C until analysis. A corticosterone ELISA kit (Enzo Life Sciences) was used to quantify corticosterone levels in blood serum samples. Samples were first brought to room temperature and, per assay procedure listed in manual, serum was analyzed for presence of corticosterone. Samples were run in duplicate alongside standards, and colorimetric product outcome was assessed using a SPECTROstar Nano microplate reader (BMG Labtech) at 405nm.

### Immunohistochemistry (IHC) and Microscopy

After 1 week of post fixation, hemisphere samples were transferred to 30% sucrose solution for cryoprotection until sunk. Brains were sectioned at 40μm using a freezing microtome (Leica), stored in serial sections for PFC in a 24 well plate. Sections were stored in anti-freeze solution (80mL 0.2 M phosphate buffer, 240mL dH2O, 240mL ethylene glycol, 240mL glycerol) at -20°C. Sections were processed for PV and 5-mC using IHC, with all washes and incubations occurring on an orbital shaker at room temperature except where noted. One day one, sections were washed 3 times for five minutes in 1x phosphate buffered saline (PBS) to remove anti-freeze solution and then incubated for one hour in permeabilization solution (1xPBS with 0.5% Triton; 0.5% PBST). Sections were washed again in 1x PBS, then incubated at 37°C in 1 N hydrochloric acid (HCl) for thirty minutes. HCl was removed, sodium tetraborate (0.1 M) was added to each well and sections were incubated at room temperature for one hour. Sections were washed three time at 5 minutes each in 1x PBS and blocked in blocking buffer (2% normal goat serum (NGS), 0.2% PBST) for one hour. Next, samples were incubated with primary antibody solutions diluted in blocking buffer including mouse anti-5-mC (1:2000; Epigentek) and rabbit anti-PV (1:1000; Novus). Sections were incubated overnight on an orbital shaker at 4°C.

On the second say, sections were washed for three time at 5 minutes. Then, with the lights off, samples were incubated in fluorescent secondary antibody solutions: goat anti-mouse Alexa Fluor 488 (1:500; Life Technologies) and goat anti-rabbit Alexa Flour 555 (1:500; Life Technologies) at room temperature for one hour. Sections were washed three times at 5 minutes each in 1x PBS, and then mounted onto glass slides. ProLong™ Gold Antifade mounting medium (Invitrogen) was used to secure glass coverslips after sections were left to dry for 2 hours. Slides were sealed with clear nail polish 24 hours later and stored in a light-tight box until imaging.

Stained sections containing PL and IL PFC were imaged at 20x using a Keyence All-in-One Fluorescent Microscope (BZ-X800; Osaka, Japan)(see Figure 2B for imaging locations). Two-three brain sections were imaged per subject. For the PFC, both the prelimbic (PL) and infralimbic (IL) PFC were imaged using Cy5 (PV) and GFP (5-mC) filters. Images were taken with the following exposure times (5-mC (1/10s); PV (1/6s)) and archived for later quantification.

Image analysis was performed using ImageJ to calculate overall PV intensity, PV cell count, intensity of 5-mC in samples, and the intensity of 5-mC colocalized with PV. For intensity measurements, overall intensity of fluorescent signal in each image was calculated to determine relative levels compared to background. Intensity serves as a semi-quantitative measure for protein amount when signal intensity is compared using the same parameters across animal and cohort, with brightness standardized across samples (Crow and Yue, 2019). ImageJ parameters were set to ensure only area, mean gray value, integrated density, and limit to threshold could be calculated. PV images for each brain image were RGB split, and intensity of PV signal was calculated using the threshold window for the red channel. Threshold level was set to ensure only the cells containing PV were highlighted (within 50 threshold units for each image) and the integrated density (IntDen) values were used for data analysis as detailed below.

PV cell count was performed by opening Cy5 channel images in ImageJ and converting the image to a black and white 8-bit. Threshold tool was used to highlight all PV containing cells and the top threshold slide ranged from 40-60 units, while the bottom was set to 255. A macro was then used to calculate the IntDen of each image (run(“Analyze Particles…, “size=400-2000 pixels, circularity=0.40-1.0, show=outlines display clear”) which was then gathered and used for data analysis as explained below.

To calculate colocalized 5-mC intensity in PV cells, 5-mC and 5-mC/PV overlay images (created using Keyence Image Analyzer) were opened in ImageJ and the cells that colocalized for 5-mC and PV were traced on the overlay image using the freehand tool. Traces in the overlay were applied to the 5-mC image and the IntDen value was calculated for each image and collated.

### DNA Extractions for ELISA and Pyrosequencing

Samples were thawed at room temperature and DNA was extracted using DNeasy Blood and Tissue Kit (Qiagen) according to the manufacturer’s directions, with some modifications: tissue was homogenized for 45 seconds using a drill attached to a pestle following addition of Qiagen tissue lysis buffer. Following addition of proteinase K, tissue was placed on a Thermomixer shaking incubator at 56°C for 4 hours before adding Qiagen lysis buffer. Following extraction, DNA was analyzed for purity (260/280>1.6) using a Nanodrop (Thermo Fisher Scientific), and samples that did not meet this purity standard were excluded from analyses.

### 5-mC ELISA

To determine global DNA methylation patterns in regions of interest, the MethylFlash Global DNA Methylation (Epigentek; specific for 5-mC) ELISA kit was used per the manufacturer’s instructions. Purified DNA (adjusted per sample to 100ng/μl) for each subject was pipetted into well plates in singlets with experimental condition varying randomly across the plate. Preliminary analyses and normalization were conducted according to manufacturer’s instructions (Alshamrani et al., 2023; Okuda et al., 2023). Percent 5-mC was detected according to optical density readings using the SPECTROstar Nano microplate reader (BMG Labtech) at 450nm. Analyses for the assay were performed by running a second order logarithmic regression to calculate the total percentage of 5-mC detected in each well.

### *Pvalb* Pyrosequencing

Purified PFC DNA samples (78 samples; two samples were excluded from original cohort of 80 due to low DNA yields) were sent to EpigenDx (Hopkinton, MA) for bisulfite modification, PCR amplification and pyrosequencing analysis. 200-500ng of purified DNA per sample was used for bisulfite modification followed by the PCR amplification. Pyrosequencing was used to determine changes to DNA methylation on the *Pvalb* gene using a sequencing-by-synthesis method to quantify CpG methylation (Delaney et al., 2015). For pyrosequencing, quantification of methylation at target CpG sites was determined based on the ratio of C to T bases following bisulfite treatment and PCR (Tost and Gut, 2007).

Primers for the *Pvalb* gene 5’ untranslated region (UTR) were created based on previous literature that identified increased methylation at two CpG sites in the rat genome that likely represent transcription factor binding sites (Fachim et al., 2016). The *Pvalb* DNA sequence analyzed was located on chromosome 7 (Bases 109786127-109786092), and the CpG sites analyzed relative to the start codon were located at site #41 and site #40 (Figure 6). The transcript length was 1267 bps (118 aa). Additional primer information is the proprietary information of EpigenDx. Pyrosequencing was conducted with a PSQ™ 96HS system (Biotage, Uppsala, Sweden). Three positive controls (low, medium, and high methylated DNA) and one negative control were used for analysis. PCR bias testing was used validate the pyrosequencing assay, where different ratios of unmethylated DNA control and *in* vitro methylated DNA were mixed, processed through bisulfite modification, PCR, and pyrosequencing to assess whether expected methylation percentages correlate with those obtained from mixing, which yielded R^2^=0.9227, showing strong correlation.

### Data Analysis

All measures collected for the OFT, IHC, ELISA and pyrosequencing experiments were initially analyzed in GraphPad Prism using three-way ANOVAs to determine overall effects of ELA, sex, and age. When there were significant main effects or interactions, these were followed up with two-way ANOVAs split by sex, as prior work by our group and others suggest that males and females often show different patterns of behavior (Ellis and Honeycutt, 2021) and we were therefore interested in uncovering how ELA influences outcomes within each sex. Significant main effects or interactions were followed up with appropriate post-hoc analyses and corrected for multiple comparisons. Outliers (over two standard deviations from the mean) were determined using Grubbs Test (GraphPad) for significant outliers and, if significant (*p*<0.05), were excluded from analyses (Berger and Kiefer, 2021). All outcomes that trended toward significance were designated as having a *p* value of less than or equal to 0.08, and these were also followed up with post-hoc tests. Results from ELISA assays were fit to a normal curve, using known concentrations of either corticosterone (pg/mL) of 5-mC (total percentage), through freely available software (MyAssays.com) prior to statistical evaluation. Correlations were assessed between PV cell counts and 5-mC intensity of PV cells by computing Pearson correlation coefficients with two-tailed analyses and a 95% confidence interval.

## RESULTS

### Behavioral

#### Center Duration

A three-way ANOVA of total time (s) in center revealed a main effect of age (*F*(1,71) = 10, *p* = 0.0022. Separate two-way ANOVAs were run for each sex, with males showing no main effects or interactions, while females showed a significant main effect of age (*F*(1,36) =7.789, *p* = 0.0084). Post-hoc Šidák’s multiple comparisons revealed a trending increase in time spent in center in P45 female ELA rats compared to P25 ELA females (*p* = 0.0682). See Figure 3A.

**Figure 3.**
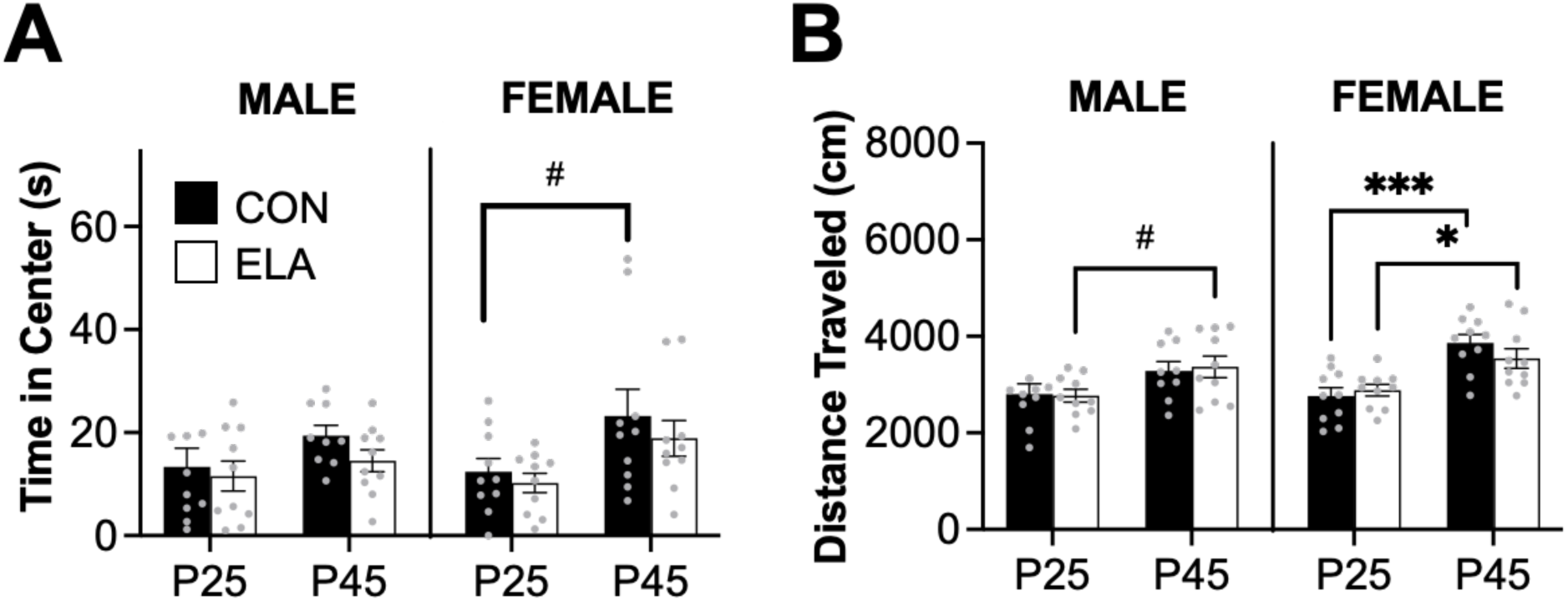
Behavior in the open field test (OFT) is driven by sex and age but is not significantly impacted by early life adversity (ELA) rearing. Indices of anxiety-like behavior assessed in the OFT did not show any significant impact of early life adversity (ELA) rearing in either time spent in center (**A**) or in distance traveled (**B**). Overall, female animals appeared more exploratory than males, especially as young adults (P45). Age significantly impacted behavior in females, with a trend toward a developmental increase in time spent in center in CON females (**A**) that was also observed for both CON and ELA females in overall distance traveled across the 10-minute test (**B**). #*p*<0.08, \**p*<0.05, \*\*\**p*<0.001

#### Distance Traveled

Three-way ANOVA on total distance traveled (cm) during the 10-minute test revealed a significant main effect of age (*F*(1,71) = 29.99, *p* < 0.0001). Two-way ANOVAs split by sex show significant main effects of age in both males (*F*(1,35) = 7.61, *p* = 0.0092) and females (*F*(1,36) = 26.55, *p* < 0.0001). Šidák’s multiple comparisons show a trending increase in distance traveled in ELA males at P45 compared to P25 (*p* = 0.0682). Multiple comparisons in females showed significant increases in distance traveled from P25 to P45 in both CON (*p* = 0.0001) and ELA (*p* = 0.019) rats. See Figure 3B.

### Corticosterone

Initial three-way ANOVA on serum corticosterone revealed significant main effects of sex (*F*(1,68) = 9.867, *p* = 0.0025) and rearing (*F*(1,68) = 10.73, *p* = 0.0017), with a significant sex x rearing interaction (*F*(1,68) = 4.529, *p* = 0.0396. There was also a trend for an age x sex interaction (*F*(1,68).= 3.274, *p* = 0.0748). A separate two-way ANOVA in males revealed no significant effects. A two-way ANOVA in females showed a significant main effect of rearing (*F*(1,35) = 9.637, *p* = 0.0038), such that P25 ELA females had significantly lower corticosterone concentrations than CON females (*p* = 0.0033) (Figure 4). In CON females, there was a trend toward a decrease in corticosterone levels from P25 to P45 (*p* = 0.0744).

**Figure 4.**
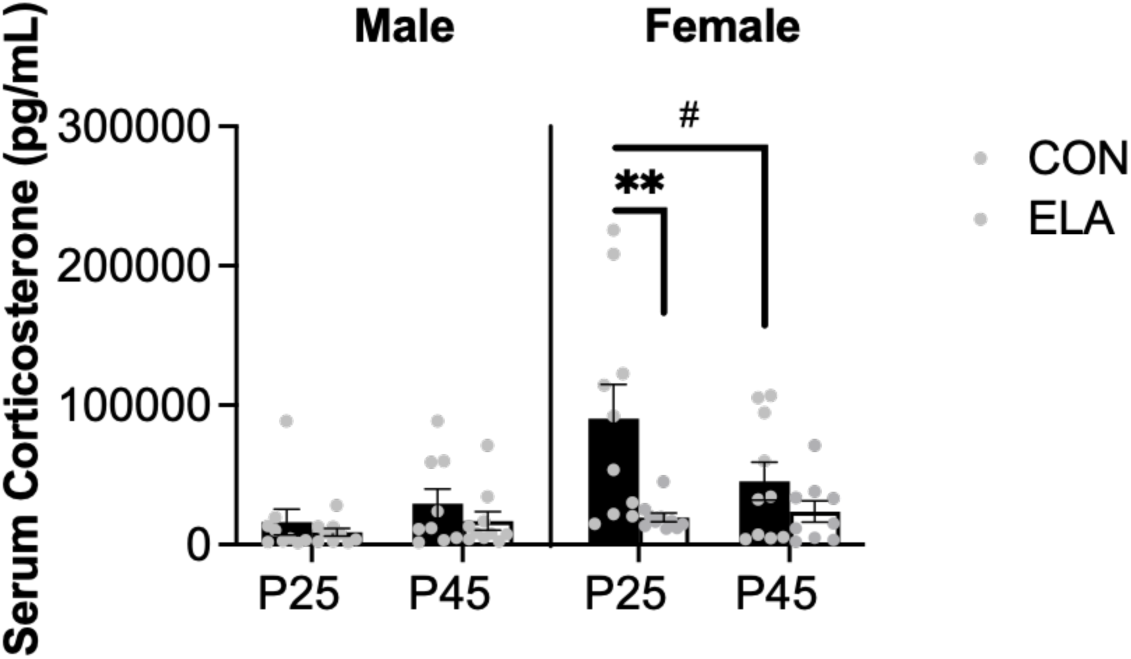
Serum corticosterone levels are reduced in female rats with a history of early life adversity (ELA). Blood serum samples collected approximately 90 minutes following behavioral assessment show sex-specific changes in serum corticosterone levels. Males do not show any significant effects of ELA or age, whereas ELA females show a significant impact of ELA in reducing serum corticosterone compared to CON females. P45 levels in ELA females are reduced compared to P25 samples, indicating an interaction between age and rearing condition on female corticosterone outcomes. **#***p*<0.08; **\*\****p*<0.01

### 5-mC ELISA for PFC Samples

ELISA assays were conducted on extracted DNA samples from PFC homogenate to determine effects of rearing, sex, and/or age on percent 5-mC methylation. Three-way ANOVA showed no significant main effects of interactions (Figure 5).

**Figure 5.**
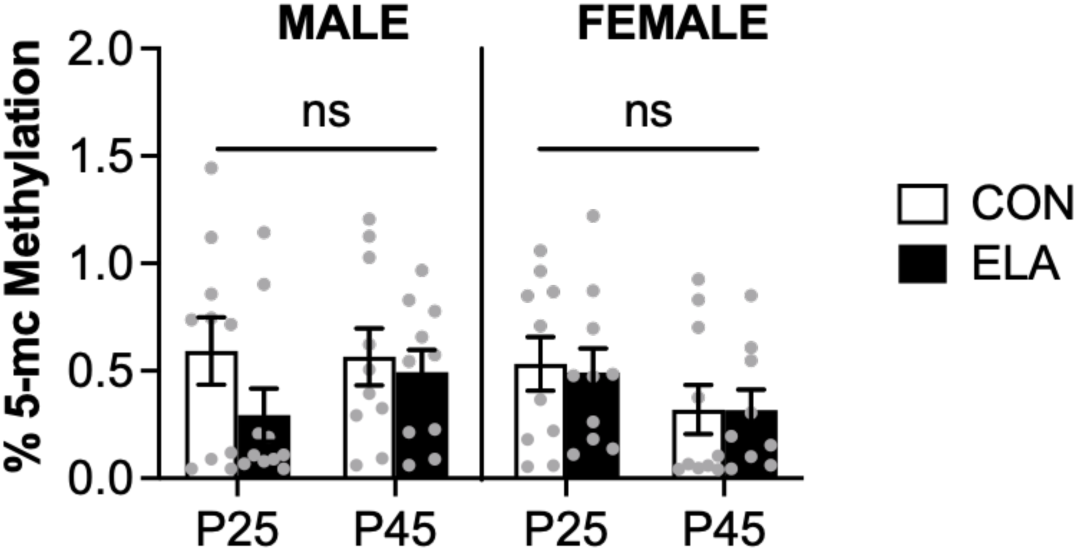
ELISA 5-mC methylation levels are not significantly impacted in a homogenous sample of the prefrontal cortex (PFC). Early life adversity (ELA) did not appear to impact overall levels of the DNA methylation marker 5-mC as assessed through ELISA colorimetric assay of purified DNA from PFC. ns (non-significant).

### Changes to Parvalbumin and 5-mC

#### Parvalbumin (PV) Cell Counts

Initial three-way ANOVA of PV cell counts in PL PFC following microscopy analyses revealed a significant main effect of sex (*F*(1,65) = 4.616, *p* = 0.0354), as well as significant sex x rearing (*F*(1,65) = 8.672, *p* = 0.0045) and age x sex x rearing interactions (*F(*1,65) = 9.603, *p* = 0.0029), with a trending age x sex interaction (*F*(1,65) = 3.784, *p* = 0.0557). Follow up two-way ANOVA in males showed a significant main effect of age (*F*(1,34) = 4.793, *p* = 0.0355) and a trending main effect of rearing (*F*(1,34) = 4.098, *p* = 0.0508). Šidák’s corrected post-hoc analyses showed that PV cell count increases with age *(p* = 0.0191), such that P45 counts were significantly higher than P25 in CON rats. Additionally, ELA significantly increases PL PV cell count at P25 compared to CON (*p* = 0.0255). A two-way ANOVA to assess female PV cell counts in the PL PFC showed a significant main effect of rearing (*F*(1,31) = 4.604, *p* = 0.0398) and a significant rearing x age interaction (*F*(1,31) = 7.254, *p* = 0.0113). Šidák’s corrected post-hoc analyses revealed a trend toward an overall decrease in PV cell count in CON female rats from P25 to P45 (*p* = 0.0503), and at P25 ELA females showed a significant decrease in PV cell count compared to CON (*p* = 0.0031). See Figure 6A.

**Figure 6.**
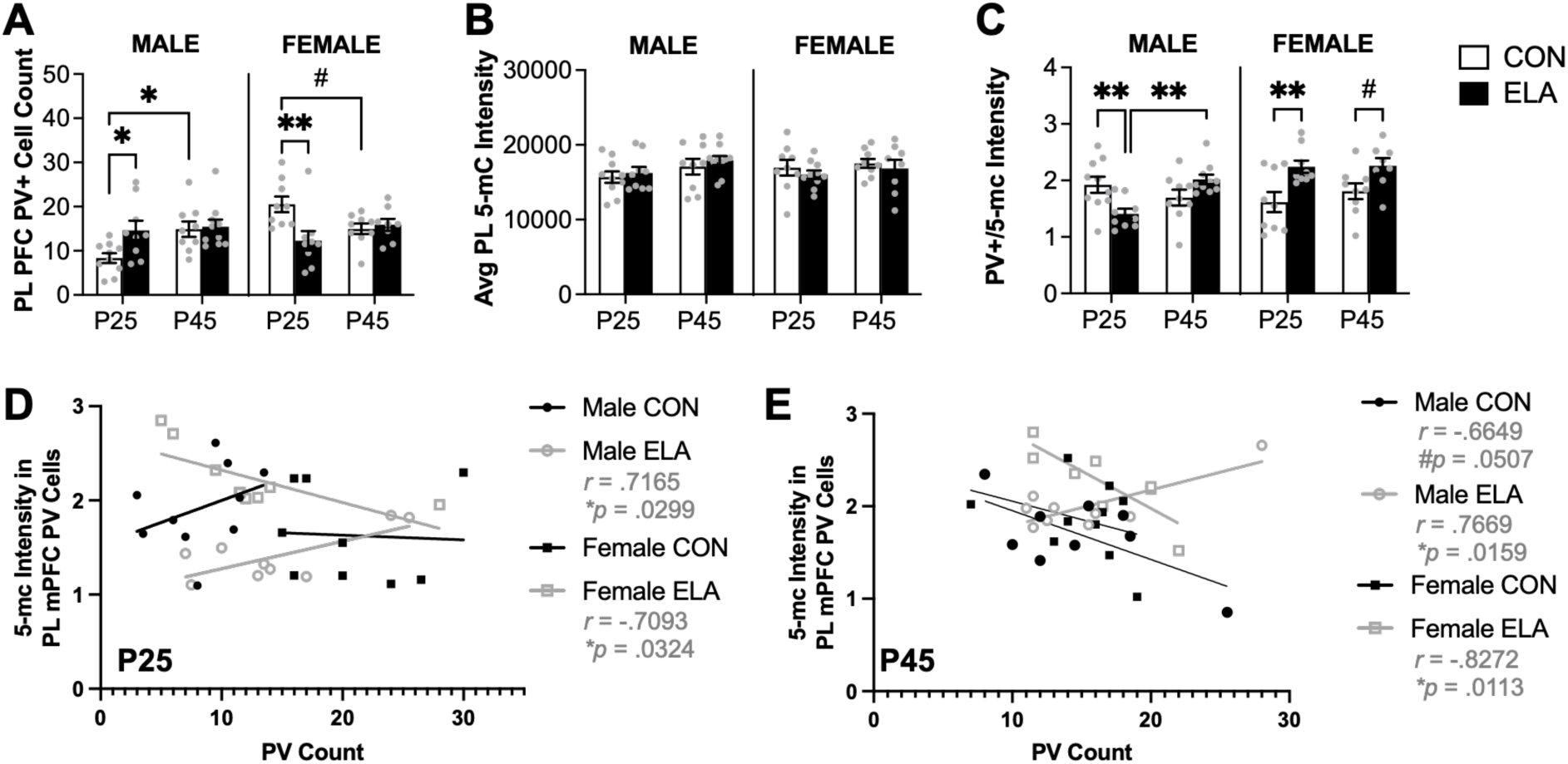
Prelimbic (PL) PFC PV cell counts and 5-mC intensity in PV cells show a sex-specific relationship following early life adversity (ELA). Within the PL PFC, males and females show opposing effects of ELA on PV cell count that is pronounced at P25 (**A**). Both males and females appear to show specific developmental patterns of PV, which is evidenced by changes in PV cell count in CON rats from P25 to P45. Robust changes in PV cell count are induced by an interaction with rearing and sex at P25, such that ELA males have increased PV cell count compared to CON males. In females, PV cell count is significantly reduced compared to female CON rats. In both sexes, ELA influences PV cell count in a way that makes juvenile (P25) ELA PV counts comparable to young adult (P45) counts. When observing overall 5-mC intensity within the PL PFC, there were no significant differences (**B**); however, this accounts for 5-mC levels across all cell types in the section. When 5-mC intensity analyses were restricted to cells that colocalized with PV, there were significant effects in both males and females (**C**). In males, 5-mC significantly intensity increases in ELA rats from P25 to P45. At P25, ELA males show a robust reduction in 5-mC intensity in PV cells compared to CON. In in P25 females, ELA serves to significantly increase 5-mC intensity colocalized with PV cells, and this pattern appears to persist into young adulthood (P45). Notably, in ELA rats PV cell counts show a strong correlation with 5-mC intensity of PV cells at both P25 and P45. Male ELA rats show a significant positive correlation, such that as levels of 5-mC intensity of PV cells increases, PV cell count also increases. In females, there is a strong negative correlation, such that as 5-mC intensity increases in PV cells, PV cell count decreases (**D**). These same relational patterns are maintained in ELA males and females into young adulthood (**E**) within the PL PFC. #*p*<0.08; \**p*<0.05, \*\**p*<0.01.

Within the IL PFC, three-way ANOVA revealed a significant sex x rearing interaction (*F*(1,65) = 5.374, *p* = 0.0236) on PV cell count. Follow up two-way ANOVA in males showed a significant main effect of rearing (*F*(1,34) = 5.699, *p* = 0.0227), with a trending effect of age (*F*(1,34) = 3.389, *p* = 0.0744); however, there were no significant post-hoc multiple comparisons. In females, a two-way ANOVA revealed no significant main effects or interactions in IL PV cell counts (Figure 7A).

**Figure 7.**
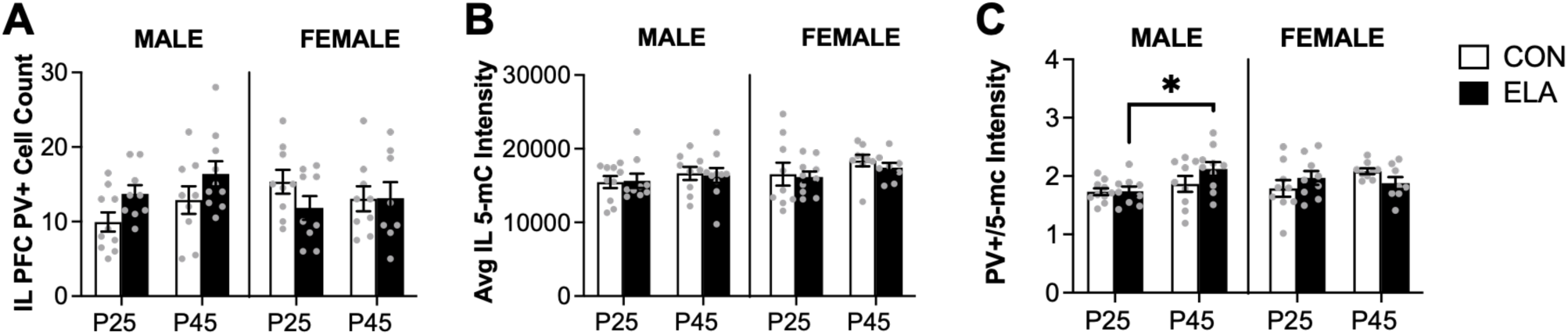
Infralimbic (IL) PFC PV cell counts and 5-mC intensity are not significantly impacted by early life adversity (ELA). ELA does not significantly change PV cell counts in the IL PFC in males or females (A). Like the PL PFC, 5-mC intensity across all cell types in IL PFC samples is not impacted by ELA in males or females (B). Interestingly, when analyses are restricted to assessing 5-mC intensity in colocalized PV cells, ELA males show an increase in intensity from P25 to P45 (**C**). **\****p*<0.05

### 5-mC Intensity

Three-way ANOVA of 5-mC intensity in the PL PFC revealed no significant main effects or interactions. However, there was a trending effect of age (*F*(1,66) = 3.63, *p* = 0.0611), which warranted follow-up two-way ANOVAs split by sex. Only males showed a trend toward an effect of age in the two-way ANOVA (*F*(1,34) = 3.616, *p* = 0.0657); however, there were no significant post-hoc effects and females did not show any main effects or interactions in PL PFC (Figure 6B).

Initial three-way ANOVA of 5-mC intensity in the IL PFC also showed no significant main effects of interactions, though did show a trending effect of age (*F*(1,66) = 3.388, *p* = 0.0702), thereby warranting follow up analyses split by sex, though two-way ANOVAs showed no significant main effect or interactions driving 5-mC intensity in males or females (Figure 7B).

### 5-mC Intensity in PV Cells

To determine whether 5-mC intensity differed in PV cells, 5-mC intensity analysis was restricted to cells that also colocalized with PV. Within the PL PFC, a three-way ANOVA evaluating 5-mC intensity of PV cells revealed significant main effects of sex (*F*(1,65) = 5.794, *p* = 0.0189) and rearing (*F*(1,65) = 5.825, *p* = 0.0186), as well as significant sex x rearing (*F*(1,65) = 11.88, *p* = 0.001)and age x sex x rearing (*F*(1,65) = 7.589, *p* = 0.0076) interactions. Follow-up two-way ANOVAs were split by sex. Two-way ANOVA in males revealed a significant rearing x age interaction (*F*(1,34) = 12.80, *p* = 0.0011), such that Šidák’s multiple comparisons show ELA males show an increase in 5-mC intensity with age (*p* = 0.0016), and at P25 ELA males have significantly decreased levels of 5-mC intensity in colocalized PV cells (*p* = 0.0076). A two-way ANOVA for females showed a significant main effect of rearing (*F*(1,31) = 14.08, *p* = 0.0007). Šidák’s post-hoc comparisons revealed that ELA females had significantly higher intensity of 5-mC in PV cells compared to CON at P25 (*p* = 0.0075), with the same pattern trending at P45 (*p* = 0.0715) in the PL PFC. See Figure 6C.

5-mC intensity in IL PFC PV cells was evaluated with a three-way ANOVA showing a significant main effect of age (*F*(1,65) = 5.816, *p* = 0.0187), and a significant age x sex x rearing interaction (*F*(1,65) = 4.435, *p* = 0.0391). Follow up two-way ANOVAs split by sex revealed a significant main effect of age in males (*F*(1,34) = 6.422, *p* = 0.016), with Šidák’s post-hoc multiple comparisons test showing a significant increase in 5-mC intensity of colocalized PV cells in ELA males from P25 to P45 (*p* = 0.024). Females did not show any main effects or interactions in the two-way ANOVA. See Figure 7C.

### Correlations between 5-mC intensity in PV cells and overall PV cell count in PL PFC

To explore the within-subjects relationship between 5-mC intensity in colocalized PV cells and overall PV cell counts, Pearson’s *r* values were calculated with a two-tailed 95% confidence interval. Correlations were conducted for each age (P25 and P45) to determine the relationship between ELA and sex at each age in the PL PFC, since this region showed robust changes in response to manipulations in both PV cell count and 5-mC intensity of PV cells. At P25 within the PL PFC, only ELA rats showed a significant correlation between these factors, such that ELA males had a positive correlation between the two variables (*r*(7) = 0.7165, *p* = 0.0299), while ELA females had a negative correlation between the two variables (*r*(7) = -0.7093, *p* = 0.0324) (Figure 6D). At P45, correlational analyses showed the same relationship in ELA rats, with ELA males showing a positive correlation (*r*(7) = 0.7669, *p* = 0.0159) and ELA females showing a negative correlation between PV count at 5-mC intensity in PV cells (*r(*6) = -0.8272, *p* = 0.0113)(Figure 6E). At P45, CON males showed a trending negative correlation between PV cell counts and 5-mC intensity in PV cells (*r*(7) = -0.6649, *p* = 0.0507).

### *Pvalb* Pyrosequencing

Two CpG sites of interest on the PV promotor, *Pvalb*, were identified based on prior work suggesting their involvement in models of affective dysfunction. Purified DNA samples from the PFC were sent to a vendor for pyrosequencing (*n*=78), to determine the percentage of methylation at these identified CpG sites within the promotor region. *Pvalb* primers were created for the 5’ untranslated region (UTR) based on previous work that identified increased methylation at two CpG sites in the rat genome that likely represent transcription factor binding sites (Fachim et al., 2016). The *Pvalb* DNA sequence analyzed was located on chromosome 7 (Bases 109786127-109786092), and the CpG sites analyzed relative to the start codon were located at site #40 and site #41. There were no significant differences in methylation percentage at CpG site 40 according to a three-way ANOVA (Figure 8A). Despite having lower over levels of methylation, a three-way ANOVA for % methylation at CpG site 41 revealed a significant main effect of age (*F*(1,70) = 9.801, *p* = 0.0025). Follow up two-way ANOVAs split by sex revealed a significant main effect of age for males (*F(*1,36) = 4.369, *p* = 0.0437), but showed no significant post-hoc comparisons. In females, a two-way ANOVA also showed a significant main effect of sex (*F*(1.34) = 5.496, *p* = 0.025), with follow up Šidák’s multiple comparisons showing a trend where P25 ELA females show higher levels of methylation compared to P45 ELA females (*p* = 0.0542) (Figure 8B).

**Figure 8.**
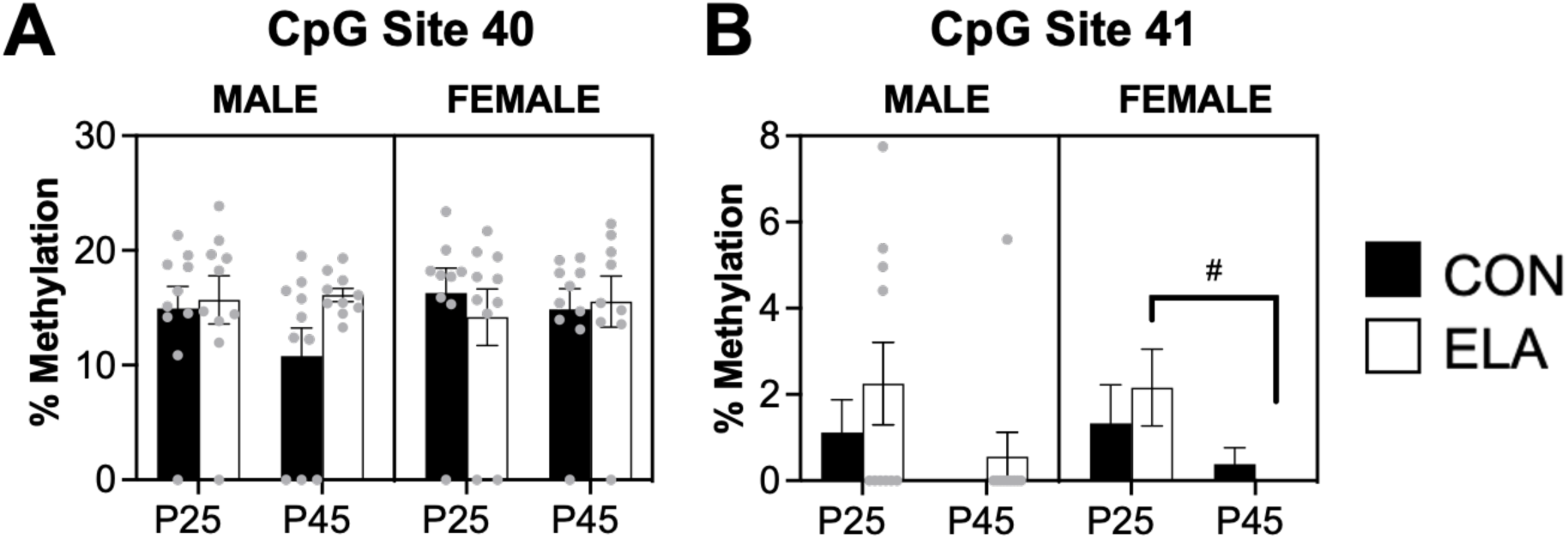
Pyrosequencing of CpG sites on the *Pvalb* promotor region show little impact of early life adversity (ELA), sex, or age on methylation outcomes. We assessed 2 CpG sites on the *Pvalb* promotor that have been previously shown to have altered methylation levels in preclinical model of psychiatric disease (Fachim et al., 2016), identified as CpG Site 40 (**A**) and CpG Site 41 (**B**). Overall, these two sites, particularly site 41, showed low levels of methylation. At site 41, there was a modest effect of age in ELA, where ELA females showed a decrease in methylation percentage when comparing levels at P25 to P45 (**B**). #*p*<0.08

## DISCUSSION

In the present work, male and female rats were exposed ELA in the form of maternal separation and anxiety-like behavior, neural, and epigenetic impacts were assessed at two developmental timepoints (juvenile, P25 and early adolescence, P45). In the PL and IL PFC PV cell count, 5-mC intensity, and 5-mC intensity colocalized with PV cells were evaluated to robust impacts of age, sex, and rearing condition. We also explored serum corticosterone levels in all conditions to determine the degree to which ELA impacts circulating stress hormone levels. Since prior work suggests that juvenile ELA exposed female rats show earlier maturation in PFC developmental profile compared to male rats (Honeycutt et al., 2020), we expected to see sex-specific effects across multiple domains. Based on past work, we also expected to see a decrease in PV cell count (Godavarthi et al., 2014; Lussier and Stevens, 2016) in the PFC following ELA, especially in juveniles with parallel increases in indices of DNA methylation (5-mC intensity). Our findings revealed robust sex-dependent changes in PV cell in count in ELA juvenile male and female rats in the PL PFC, with only minimal impact observed in the IL PFC. Specifically, PV cell count was decreased in PL PFC in juvenile ELA females compared to CON, with juvenile ELA levels comparable to young adult cell counts. The opposite was observed in males, with ELA males showing an increase in PV cell counts compared to CON, with ELA juveniles showing cell counts comparable to young adult males. A potential linkage between lowered PV expression and PV cell count is that less PV could create calcium homeostasis imbalance, resulting in PV cell death (Godavarthi et al., 2014). However, given that males showed an ELA-related increase this may not be the case, or there may be additional factors interacting with ELA that are sex-specific.

We next sought to understand *how* PV cell counts are altered in the PFC of ELA-exposed rats and looked to epigenetics to explain these differences. First, we explored global methylation patterns (both through extracted DNA in an ELISA, as well as through 5-mC intensity measures via immunostaining) to determine if there were alterations in 5-mC methylation percentage and/or intensity following ELA within the PFC. Globally, there were no differences in 5-mC percentage regardless of rearing condition, developmental age, or sex in either approach. However, these approaches only consider gross methylation across a homogenous cell population of an entire region and cannot discern methylation differences that may be present at the neuronal subtype or gene/promotor level. Thus, 5-mC intensity was next analyzed in terms of colocalization intensity with PV cells in PL and IL PFC. Since the *Pvalb* promoter is susceptible to methylation (Fachim et al., 2016), and DNA methylation can cause a decrease in transcription of a gene (Szyf, 2013b), we expected to see an increase in DNA methylation intensity in PV cells (Houtepen et al. 2016), and an increase in DNA methylation on the *Pvalb* promoter (Fachim et al., 2016) in the PFC following ELA.

We conducted intensity analyses of 5-mC specific for PV colocalized cells. Results paralleled those observed in PV cell counts, with differential effects based on sex, age, and rearing. Juvenile ELA males showed a significant decrease in 5-mC intensity in colocalized PV cells compared to CON; whereas juvenile ELA females had a significant increase in 5-mC intensity in PV cells in the PL PFC when compared to control females of the same age, a difference that also extended, although modestly, into young adulthood. This suggests that ELA induces alterations in DNA methylation levels following ELA that appear to be specific to PV cells. Interestingly, PV count analyses show that females exposed to ELA have significantly lowered PFC PV in juvenility. By late adolescence, however, both CON and ELA animals have similar PV cell counts. PV cell count has been shown to subtly decrease in the PFC throughout development, due to neural pruning of synapses (Ellis and Honeycutt, 2021). Since females exposed to ELA already show decreases in PV cell count in juvenility, this suggests that ELA may shunt PV development in a sex-specific fashion, resulting in juvenile females with significantly less PV cells before stages of neural pruning. The early maturation of female PV that is comparable to the post-pruning stage may be indicative of dysregulation of E:I balance in the PFC, an effect that is in line with the earlier maturation of female PFC-BLA circuitry seen following ELA (Honeycutt et al., 2020).

PV expression also decreases in juvenility for ELA females, but not males (Holland et al., 2014). Importantly, when PV cell counts and 5-mC intensity in colocalized PV cells was assessed, both male and female ELA rats showed significant relationships between these two factors that persisted from juvenility to young adulthood. Specifically, in ELA males, as PV/5-Mc intensity levels increased, PV cell count increased; whereas, in ELA females, as PV/5-mC intensity increased, PV cell count decreased. Based on these results, it is possible that 5-mC accumulation may differentially mediate PV expression as a function of sex and that DNA methylation may act on the PV promoter to reduce the expression of PV, resulting in higher 5-mC intensity and lower *Pvalb* expression. Pyrosequencing and analysis of two CpG sites (CpG site 40 and 41) on the *Pvalb* promoter was conducted to determine whether PV outcomes may be linked to specific CpG site methylation on the promotor. These sites were chosen based on previous literature, and likely represent transcription factor binding sites (Fachim et al., 2016). This analysis presented low methylation levels on CpG site 41 (<5%), indicating that methylation at this site likely does not contribute significantly to differential expression of the *Pvalb* gene. CpG site 40 showed higher methylation levels (>15%), though there were no significant effects observed from our manipulations on methylation percentage at this site. These findings are inconsistent with past findings of hypermethylation of CpG site 40 for both MDD patients (Thaweethee-Sukjai et al., 2019) and schizophrenia rodent models (Fachim et al., 2016). However, the fact that there was a lot of variability in samples and many samples that showed no methylation, this may obscure subtle effects. Nonetheless, CpG site 40 and 41 are not the only CpG sites present within the *Pvalb* promoter, and both CpG 40 and 41 are ∼1600 bps from the TSS. Since previous literature describing methylation patterns on the *Pvalb* promoter is limited, methylation at other unidentified CpG sites within this region may also contribute to sex-specific alterations of the *Pvalb* gene.

While DNA methylation at CpG site 40 and 41 on the *Pvalb* promoter likely does not significantly contribute to regulation of *Pvalb* transcription, PV levels are found to be dysregulated in juvenile male and female rats exposed to ELA. This could be due to DNA methylation at other CpG sites on the *Pvalb* promoter. However, it is important to note that these approaches use heterogenous cell populations to assess changes to 5-mC as it relates to PV. Due to the physiological properties of PV and it’s specific disruption following ELA, it is possible that methylation alterations at the *Pvalb* promotor may only be appreciated in heterogenous cell populations. Alternatively, other factors, such as dopamine levels, could play a role in this dysregulation. Dopamine is a neurotransmitter that can increase *Pvalb* expression in the mPFC of rodents during development, suggesting that dopaminergic innervation may act on PV cells to accelerate the rate of *Pvalb* expression (Porter et al., 1999). Additionally, the dopaminergic system is involved in anxiety, with dopamine receptor D2 playing a critical role in dampening anxiety response (Zarrindast and Khakpai, 2015). Thus, modulation of dopamine levels could contribute to differential *Pvalb* expression in animals exposed to ELA.

Moreover, mouse models of reversible PV cell inhibition demonstrate that PV activity is required during juvenility for proper maturation of the PFC and, in mice with inhibited PV function, there are consistent alterations in behavior (Canetta et al., 2022). Another potential mediator of PV circuitry is via changes in Brain Derived Neurotrophic Factor (BDNF). BDNF is a neurotrophic factor that induces perineuronal net (PNN) formation (O’Connor et al., 2019). PNNs surround PV cells, and help stabilize surrounding neuronal networks, functionally helping inhibit the rewiring of existing circuitry (Gildawie et al., 2020; Tanti et al., 2022). Thus, variation in BDNF level may alter the stability of PNNs in the short-term and affect PV expression long-term (O’Connor et al., 2019). Overall, there are many potential contributors to changes in PV expression following ELA outside of epigenetics, though we provide compelling correlational finings here that link altered 5-mC intensity in colocalized PV cells with overall PV cell count.

Following ELA, we expected to see an increase in corticosterone reactivity for males and a decrease in corticosterone reactivity for females (Kalinichev et al., 2002; Voellmin et al., 2015). Indeed, juvenile ELA females displayed less corticosterone reactivity compared to CON animals at the same age, suggesting that ELA can blunt corticosterone reactivity. However, male rats displayed no significant changes to corticosterone reactivity regardless of rearing or developmental conditions. The match/mismatch hypothesis of psychiatric disorders may be used to in part explain why ELA may result in blunted levels of corticosterone specific for juvenile female animals (Schmidt, 2011). Moreover, coping mechanisms, which may be behavioral or physiological, associated with ELA can shape reactions to stressors later in life (Schmidt, 2011), and female rats may display blunted corticosterone due to abnormally elevated levels of corticosterone during ELA. While blunted stress hormone response could be characterized as better from a homeostatic viewpoint, impaired cortisol reactivity (Nijm et al., 2007) and lowered diurnal stress hormone levels (Chrousos and Gold, 1992) can be a precursor for the development of adverse health conditions (Chrousos and Gold, 1992; Nijm et al., 2007).

Finally, we expected to see an increase in anxiety-like behavior following ELA, traditionally characterized as a decrease in time spent in the center of the open zone of the OFT (Wigger and Neumann, 1999). However, while increased time spent in the center area for the OFT has traditionally signified a decrease in anxiety-like behavior, these results are inconsistent, especially when considering sex-differences (Ellis and Honeycutt, 2021). Previous literature suggests that female rats exhibit higher levels of locomotor activity than male rats, regardless of rearing condition (Börchers et al., 2022), which may predispose female rats to travel further distances, and enter the center zone more times due to hyperactivity compared to their male counterparts, rather than anxiety-like phenotypes. While CON and ELA rats in our experiment showed no significant changes in time spent in center or distance travelled when looking at the two developmental stages in isolation, over time the two rearing conditions display changes in locomotor activity. Specifically, ELA and CON females both show a developmental increase in exploration in the OFT.

## Limitations

The relationship between experience, genes and affective outcomes has been a puzzle for centuries. Researchers have attempted to characterize how the dynamic interplay between nurture and nature may result in a psychiatric disorder for one individual, but not another. Thus, while the results we present here suggest a sex-dependent relationship between DNA methylation and PV-specific outcomes following ELA, our methods cannot show a cell-specific impact of *Pvalb* methylation in a homogenous cell population. However, our results still present important insight into potential windows for understanding susceptibility to psychiatric disorders through epigenetic mechanisms.

Due to the psychiatric manifestations of ELA, we need pre-clinical models of ELA to understand the potential impacts of DNA methylation at certain loci on these outcomes, so that exposure to adversity may be systematically manipulated to confirm a causal relationship. To better understand the biochemical mechanisms behind psychiatric disorders and improve outcomes for treatment resistant patients, it is important to utilize animal modeling as a translational research-based approach (Phillips and Roth, 2019). Additionally, both behavioral and neurological testing is important to convey the relationship between alterations in the brain and corresponding behavioral manifestations. There are caveats to exploring animal models, including an inability to entirely attribute the anxiety-like behavior of rodents to a clinical representation of anxiety in humans. Additionally, it is difficult to determine causality, in that both environment and genetic predispositions, as well as individual variation, contribute immensely to pathological outcome, such that one source alone cannot be fully attributed to the result. Thus, the changes reported here are correlational, in that we cannot entirely exclude the possibility that the findings result from reasons other than ELA rearing, sex, or developmental time point.

One limiting factor in utilizing rodent research is that rodent models most often directly explore DNA methylation patterns related to psychiatric disease in brain tissue, while human studies utilize more indirect peripheral blood and saliva samples. Fortunately, saliva and blood samples are accurate predictors of brain DNA methylation patterns (Braun et al., 2019). Thus, even without access to brain tissue methylation levels we can have some degree of confidence that DNA methylation analyses in rodent ELA studies can be accurately leveraged to understand human outcomes.

## Conclusions

Diagnoses of psychiatric disorders, including anxiety, have skyrocketed in recent decades, and have become especially prevalent following the COVID-19 pandemic. Moreover, children exposed to ELA have an increased likelihood of developing anxiety disorders later in life. The relationship between genes and the environment may help reveal an individuals’ susceptibility to developing an affective disorder, but it is difficult to draw causal connections between neurobiological underpinnings and psychiatric outcomes. Nonetheless, it is important to leverage animal models to characterize anxiety-like phenotypes and to explore their underlying neurobiological and epigenomic drivers, as this characterization is crucial to eventually understanding how ELA may rewire the brain to increase susceptibility to mood disorders. This study contributes to that understanding, by exploring the relationship between ELA, anxiety-like behaviors, stress response, and neural markers indicative of epigenetic changes. In patients with anxiety disorders, there is a dysregulation of the E:I balance in the mPFC, such that excitation cannot be properly dampened. Additionally, ELA is known to alter the epigenomic landscape, though until now, there have been no studies directly aiming to link ELA associated changes in behavior and PV outcomes to PV-specific DNA methylation. The present findings detail sex-specific changes in PV levels in the PL PFC that are paralleled by correlated changes in 5-mC DNA methylation, suggesting that changes in DNA methylation could be responsible for ELA-induced PV pathology. Future work should aim to harness heterogenous samples to assess additional CpG sites on the *Pvalb* promoter that are closer to the transcriptional start site (TSS), and that may more directly contribute to *Pvalb* expression.

## Acknowledgements

The authors would like to acknowledge Bowdoin student colleagues who were instrumental in their support and in collecting the data presented here, including: Sydney Bonauto, Seneca Ellis, Khushi Patel, and Lucy O’Sullivan. We are grateful to our laboratory technician, Elizabeth Mann, as well as Dr. Anja Forche, for all their help and feedback with experiments and protocols. We would also like to thank Dr. Danielle Dube and Dr. Anne McBride for serving as readers and providing critical feedback on initial versions of this manuscript. Finally, we would like to thank Dr. Hai-Ying Shen and colleagues for kindly sharing the 5-mC IHC protocol with us.

## Data Availability

Data is available and can be obtained at this time by contacting the corresponding author. Data is in the process of being submitted to a repository to be made openly available upon publication of this work.

## Author Contributions

ESN and AC contributed to drafting the manuscript, collected, and analyzed data, and assisted in data interpretation. YP collected and analyzed data and provided feedback on the manuscript. JAH developed the research plan and designed experiments, drafted the manuscript, and provided training, research oversight, and data analysis and interpretation. All authors contributed to editing and reviewing the manuscript.

